# Directional Matching of Swimming Polarity Provides a Competitive Advantage During Bacterial Magneto-Aerotaxis

**DOI:** 10.64898/2026.01.09.698624

**Authors:** Carina Weigel, Daniel Pfeiffer

## Abstract

**Background:** Magnetotactic bacteria (MTB) utilize magnetosomes to align passively with Earth’s magnetic field. Magnetic alignment, coupled with flagellar motility and aerotaxis, enables MTB to perform magneto-aerotaxis—a strategy that constrains their movement to a one-dimensional trajectory along geomagnetic field lines, optimizing their search for low-oxygen niches in aquatic environments. Beyond axially constrained movement, environmental MTB isolates exhibit a hemispherically determined swimming polarity—favoring either magnetic north or south—that has been suggested to facilitate descent into oxygen-depleted zones. However, a direct quantitative evaluation of how matching swimming polarity influences navigation toward low-oxygen environments has remained elusive. Here, we employed microcapillary assays to assess the functional significance of polar magneto-aerotaxis in the model organism *Magnetospirillum gryphiswaldense*.

**Results:** We found that a magnetic field configuration matching the predominant swimming polarity of the population results in an up to 4-fold increased peak intensity of the aerotactic band compared to populations with non-matching polarity. Competition assays using fluorescently labeled north- and south-seeking populations confirmed that congruence between swimming polarity and magnetic field orientation markedly improves aerotactic band formation in oxygen gradients. Alongside our main findings, we noted biomagnetism-independent phototactic responses integrated with aerotaxis, driving collective unidirectional migration along the oxygen gradient.

**Conclusions:** Our results provide direct evidence that matching swimming polarity with the magnetic field confers a clear advantage in navigating oxygen gradients. These findings reinforce the role of the geomagnetic field in shaping MTB behavior and highlight the adaptive value of magnetotactic swimming polarity in environmental navigation.

## Background

Organisms across diverse life forms are known to perceive and respond to the Earth’s magnetic field [1, 2]. The most ancient and experimentally accessible magnetosensitive organisms are magnetotactic bacteria (MTB) [3, 4]. They orient along geomagnetic field lines using specialized intracellular organelles called magnetosomes [1–3]. Passive magnetic alignment is complemented by flagella-driven motility and chemotactic responses—most notably aerotaxis, which plays a dominant role by guiding movement toward favorable oxygen concentrations [4]. This combined strategy was termed magneto-aerotaxis, emphasizing a magnetically assisted form of aerotaxis [5–7]. It is assumed that by constraining movement along geomagnetic field lines, MTB reduce a three-dimensional random walk to a one-dimensional search, thereby enhancing their ability to locate micro- or anoxic niches that support growth and magnetosome formation [4, 5, 7]. This is reflected in the characteristic aerotactic band formation by cultured MTB in oxygen gradients, aligning with their microaerophilic lifestyle [5–9].

Beyond geomagnetic alignment, MTB populations from natural environments typically show a directional preference for one magnetic pole—referred to as “polar magneto-aerotaxis” or “swimming polarity” [1, 5–8, 10]. Polar magneto-aerotaxis is commonly investigated using the “hanging drop assay,” where a droplet of cell suspension is placed on an inverted coverslip positioned on an O-ring adjacent to a bar magnet and exposed to air [5, 6]. Using this assay, MTB from the Northern and Southern Hemispheres were found to exhibit north- and south-seeking swimming polarities, respectively, as indicated by their predominant accumulation at the edge of the drop facing the south or north pole of the magnet [6]. This directional behavior aligns with current Earth’s magnetic field orientation (where the North Magnetic Pole corresponds to the south pole of a physical magnet) and likely facilitates a downward-directed response into less oxygenated or anoxic regions in both hemispheres, away from zones with potentially detrimental oxygen levels [6]. In natural MTB populations, upward geomagnetic field inclinations greater than 6° were associated with an enrichment of south-seeking cells over north-seeking cells by more than an order of magnitude, suggesting that polar magneto-aerotaxis offers an advantage even at relatively shallow inclinations [11, 12]. In contrast, MTB populations from samples collected near the magnetic equator exhibited approximately equal proportions of north- and south-seeking cells [1, 11–14]. Although the mechanisms and relevance of magneto-aerotaxis in environments with near-zero inclination are unclear, these findings indicate that the vertical component of the Earth’s magnetic field plays a key role in selecting the predominant swimming polarity in natural habitats [12, 13]. This conclusion is further supported by laboratory experiments, where sediment samples were exposed to magnetic fields opposing the geomagnetic polarity of their original sampling sites, leading to an inversion of swimming polarity within several weeks [1, 12]. By contrast, the selective mechanisms and ecological significance of swimming polarity have been challenged by occasional observations of both north- and south-seeking MTB coexisting within the Northern and Southern Hemispheres [15, 16].

Studies on axenic cultures of magnetotactic cocci support environmental assessments of swimming polarity. These studies demonstrated that the magnetic field orientation influences individual polarity fractions and the overall abundance of cells during cultivation in oxygen gradient media [10], as well as the ability of polar cells to form aerotactic bands in microcapillaries [5, 6, 17]. Along similar lines, in the laboratory model *Magnetospirillum gryphiswaldense*, one of the few genetically tractable MTB, swimming polarity is closely linked to aerotaxis [7]. Strikingly, in *M. gryphiswaldense*, north- or south-seeking swimming polarity was rapidly reestablished within a few generations when magnetic fields were applied parallel or antiparallel to the oxygen gradient, respectively, but was lost when cultures were grown under uniform oxygen conditions with agitation, resulting in equal proportions of both cell types in the hanging drop assay [7]. These observations suggest that the previously reported “axial” magneto-aerotaxis—the apparent lack of swimming polarity in *Magnetospirilla* [5, 17]—might be an artifact of laboratory cultivation [7]. In addition to its selection during growth, the polar response of *M. gryphiswaldense* was inverted when the hanging drop assay was performed under anaerobic conditions [7]. Upon oxygen exposure, cells immediately swam to the opposite side of the drop (and thus to the opposite magnetic pole), suggesting that their immediate directional bias is determined by whether they experience oxygen levels above or below their optimal range [7]. Further evidence for the relevance of magnetic field polarity in navigation by *M. gryphiswaldense* was provided by microcapillary assay studies, which demonstrated that the orientation of the magnetic field relative to the oxygen gradient affects both aerotactic band formation efficiency and band stability [8, 9, 18]. In cell populations derived from oxygen gradient media, inverting the magnetic field resulted in an immediate, partial dispersal of the band, suggesting that polar subpopulations were directed toward regions above and below their preferred oxygen concentration [8].

While the studies discussed above suggest that matching swimming polarity is crucial for magneto-aerotaxis, a quantitative assessment of its role in navigating toward low-oxygen environments has not yet been performed. To address this gap, we performed microcapillary assays on polar *M. gryphiswaldense* populations, complemented by quantitative analyses and competition assays between north- and south-seeking cells. Prior to microcapillary experiments, the polarity bias of cell populations was assessed using the hanging drop assay, allowing us to directly link observations across both assays. Thereby, we found that the relative abundance of north- and south-seeking cells in the hanging drop assay is mirrored in their distribution within aerotactic bands formed under magnetic fields aligned parallel or antiparallel to the O_2_ gradient. Importantly, we demonstrate that swimming polarity aligned with the magnetic field orientation substantially enhances migration toward microoxic zones and confers a competitive advantage during magneto-aerotaxis. Moreover, we confirm that swimming polarity is selected within just a few generations based on the orientation of the magnetic field relative to the oxygen gradient. Additionally, we identify light-dependent motility responses that drive collective, unidirectional migration along the oxygen gradient. Finally, we discuss our findings not only within the context of existing literature, but also through illustrative models exploring potential mechanisms underlying swimming polarity determination and selection.

## Results

### Swimming Polarity Is Determined by Magnetic Field Orientation and Intensity

Prior to analyzing swimming polarity using the capillary assay, we aimed to obtain cultures enriched in either north- or south-seeking cells through a cultivation-dependent approach described previously [7]. Therefore, cultivation was performed in non-agitated culture tubes (to establish a vertical oxygen gradient), superimposed with a homogeneous 0.6 mT magnetic field produced by coils oriented either parallel (northern field, NF) or antiparallel (southern field, SF) to the oxygen gradient, simulating Northern and Southern-Hemisphere geomagnetic field polarity, respectively.

Consistent with prior results [7], swimming polarity typically developed after ∼10 generations of incubation. Hanging drop assays demonstrated a strong enrichment of the respective polarity type; however, cultures never contained exclusively one type (**Fig. 1A,B**). In addition to artificially generated vertical magnetic fields, cultures were incubated in the local geomagnetic field (GMF) and in a zero field (ZF) with the geomagnetic field cancelled. Although previous studies indicate that magnetospirilla exhibit rather poor alignment in physiologically relevant magnetic fields [19, 20], incubation in the GMF (see **Methods** for field parameters) already led to an enrichment of north-seeking cells relative to south-seeking cells, suggesting that the GMF is sufficient for polarity selection (**Fig. 1C**). Nevertheless, enrichment for one polarity type was notably lower compared to artificially imposed magnetic fields. As expected, incubation in a ZF environment did not promote enrichment of one polarity type (**Fig. 1D**).

**Fig. 1:**
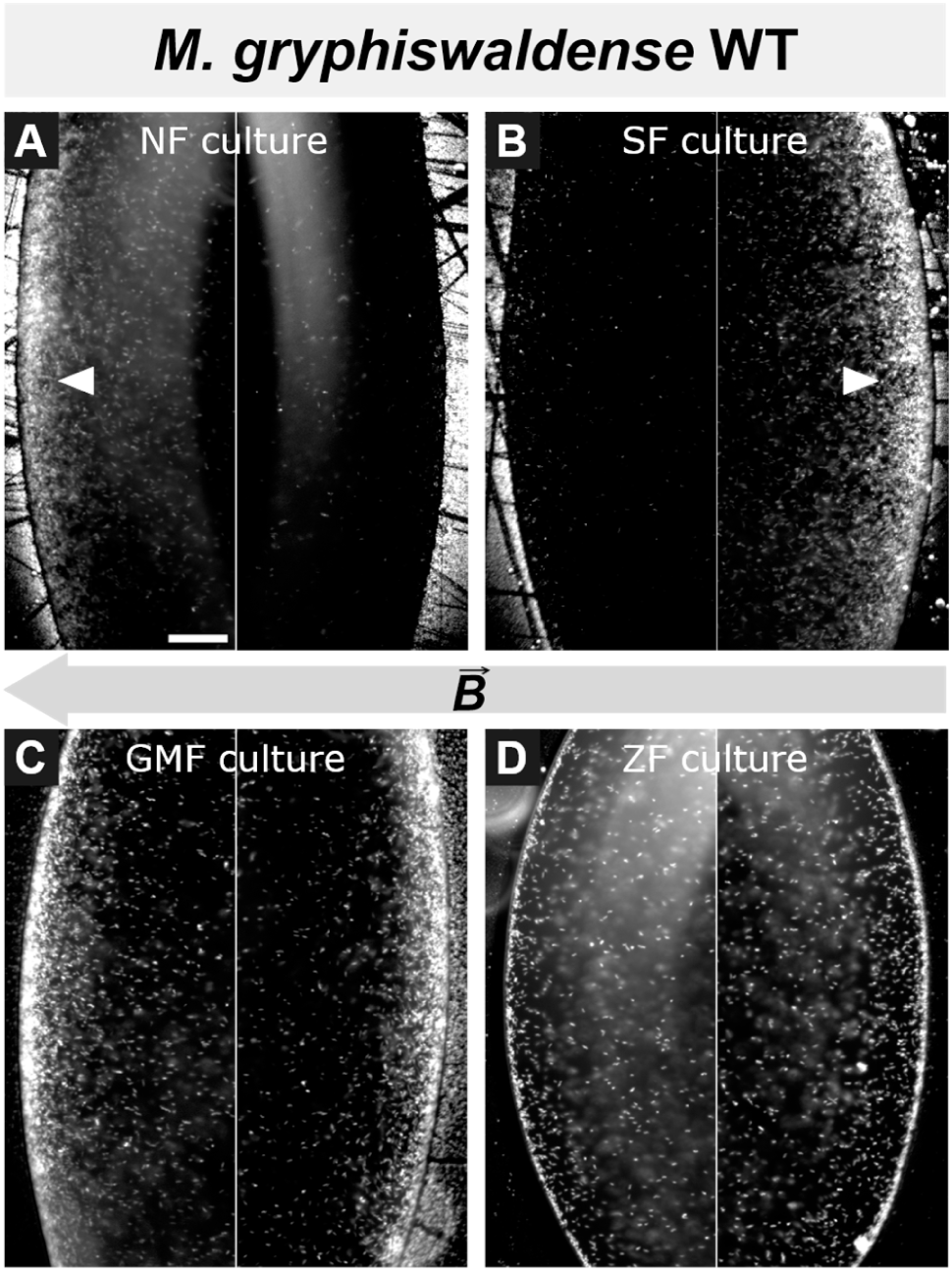
Cultivation-dependent selection of swimming polarity in *M. gryphiswaldense* wild-type cells. Swimming polarity was analyzed using hanging-drop assays. Cells accumulating at the drop edges corresponding to the north and south magnetic poles are classified as south-seeking and north-seeking, respectively. Representative images of both edges of the same drop are shown for each sample. Cultures were grown for at least 10 generations in vertical oxygen gradients under the following magnetic field configurations: **(A)** NF: uniform magnetic field applied parallel to the oxygen gradient, enriching north-seeking cells (white arrow). **(B)** SF: uniform magnetic field applied antiparallel to the oxygen gradient, enriching south-seeking cells (white arrow). **(C)** GMF: growth under the ambient geomagnetic field (Bayreuth, Germany). **(D)** ZF: ambient magnetic fields were eliminated. Arrows indicate magnetic field (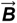) direction. Scale bar: 50 µm (applies to all images; shown in A).

### Matching Swimming Polarity Improves Navigation to Microoxic Conditions

To identify the benefits of swimming polarity on magneto-aerotaxis, culture samples were transferred into flat glass capillaries. Two capillaries, each filled with the sample and sealed at one end, were inverted relative to each other (**Fig. S1**) and monitored in parallel to observe aerotactic band formation under magnetic fields resembling Northern- and Southern-Hemisphere geomagnetic field polarity, hereafter referred to as NF and SF settings, respectively. As assessed by dark-field micrographs acquired under identical illumination, the distribution of cells in the aerotactic band under the two magnetic field polarities (**Fig. 2**; see also **Fig. S2** for overlaid intensity distributions) reflected the proportions of north- and south-seeking cells observed in the hanging-drop assay (**Fig. 1**). However, while quantification in the hanging-drop assay is limited by the curved geometry of the drop, rectangular flat glass capillaries enabled precise measurement of aerotactic band characteristics, including distance from the meniscus, band width, and band peak intensity.

**Fig. 2:**
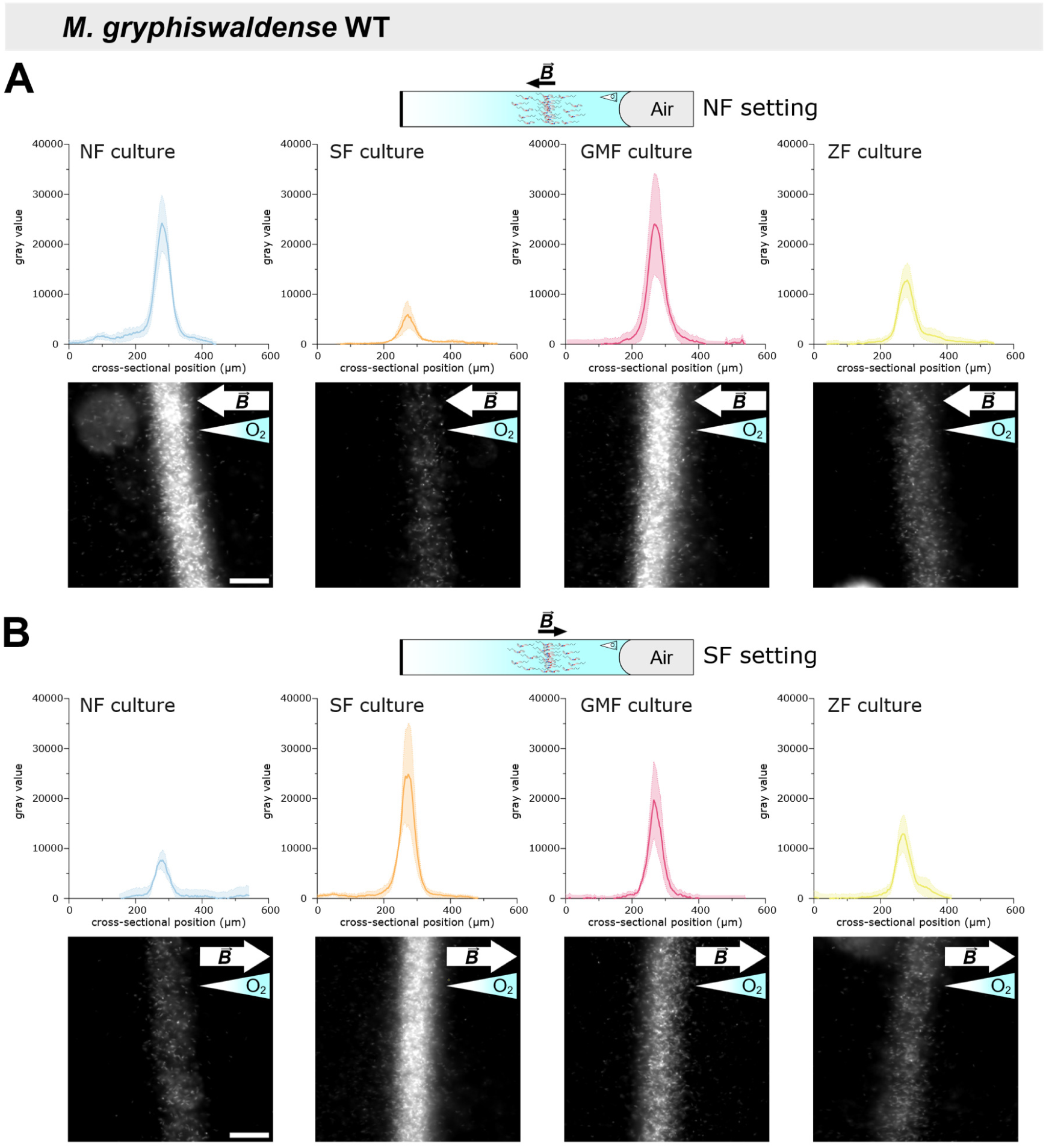
Intensity profile analysis of aerotactic bands of *M. gryphiswaldense* wild-type cultures. To study aerotactic band formation, cultures were adjusted to the same optical density and loaded into glass microcapillaries. Capillaries were sealed at one end to establish a unidirectional oxygen gradient and examined by dark-field microscopy under both NF **(A)** and SF **(B)** settings. NF and SF cultures were precultured in uniform magnetic fields oriented parallel and antiparallel to the oxygen gradient, respectively. GMF cultures were incubated in the ambient geomagnetic field, ZF cultures in zero field. Aerotactic band intensity profiles and dark-field micrographs shown correspond to the 180-min time point, when aerotactic bands had stabilized; imaging settings were kept constant across experiments. Arrows indicate magnetic field (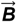) direction; gradient-filled triangles indicate oxygen gradient direction. Peak intensity (from intensity profiles) was used as a proxy for the number of cells within the aerotactic band. Intensity profiles represent the mean ± standard error of the mean (SEM) from *n* = 3 independent experiments. Scale bars: 50 µm (apply to all images; shown only in the leftmost images).

Analysis of aerotactic band intensity profiles revealed that, under the NF setting, SF cultures exhibited a 75% reduction in peak band intensity compared to NF cultures (**Fig. 2A**), indicating substantially reduced accumulation of south-seeking cells. Conversely, under the SF setting, NF cultures exhibited a marked decrease in band intensity, with the peak reduced by ∼70% relative to SF cultures (**Fig. 2B**). Furthermore, intensity profile analysis confirmed that growth under relatively weak geomagnetic field conditions (GMF cultures) was sufficient to select for north-seeking cells, as the peak aerotactic band intensity under the NF setting reached ∼99% of that observed for NF cultures (**Fig. 2A**). In contrast, under the SF setting, GMF cultures reached nearly 80% of the peak band intensity of SF cultures (**Fig. 2B**), indicating that they still contained a substantial number of south-seeking cells (consistent with hanging-drop results, **Fig. 1**) and that the artificially imposed field on NF cultures exerted stronger selective pressure. A balanced coexistence of north- and south-seeking cells in ZF cultures, as seen in the hanging-drop assay (**Fig. 1**), was reflected by aerotactic band peak intensities of ∼50% relative to NF and SF cultures under both NF and SF settings (**Fig. 2A,B**).

The distance of the aerotactic band to the air-liquid interface was monitored throughout the capillary assay, with no notable differences in dynamics observed between the four culture conditions (**Fig. S3**). Moreover, after 180 min in the microcapillary, the distance of the aerotactic band from the meniscus did not significantly differ among cultures (**Fig. 3A**). However, cells from NF cultures accumulated slightly closer to the meniscus under the NF setting, while SF-culture cells displayed the same behavior under the SF setting. This observation may reflect accelerated oxygen depletion due to higher cell density, which could lead to faster migration of the aerotactic band toward the meniscus. Finally, measurements of the full width at half maximum of aerotactic bands revealed no significant differences among the culture conditions (**Fig. S4A**).

**Fig. 3:**
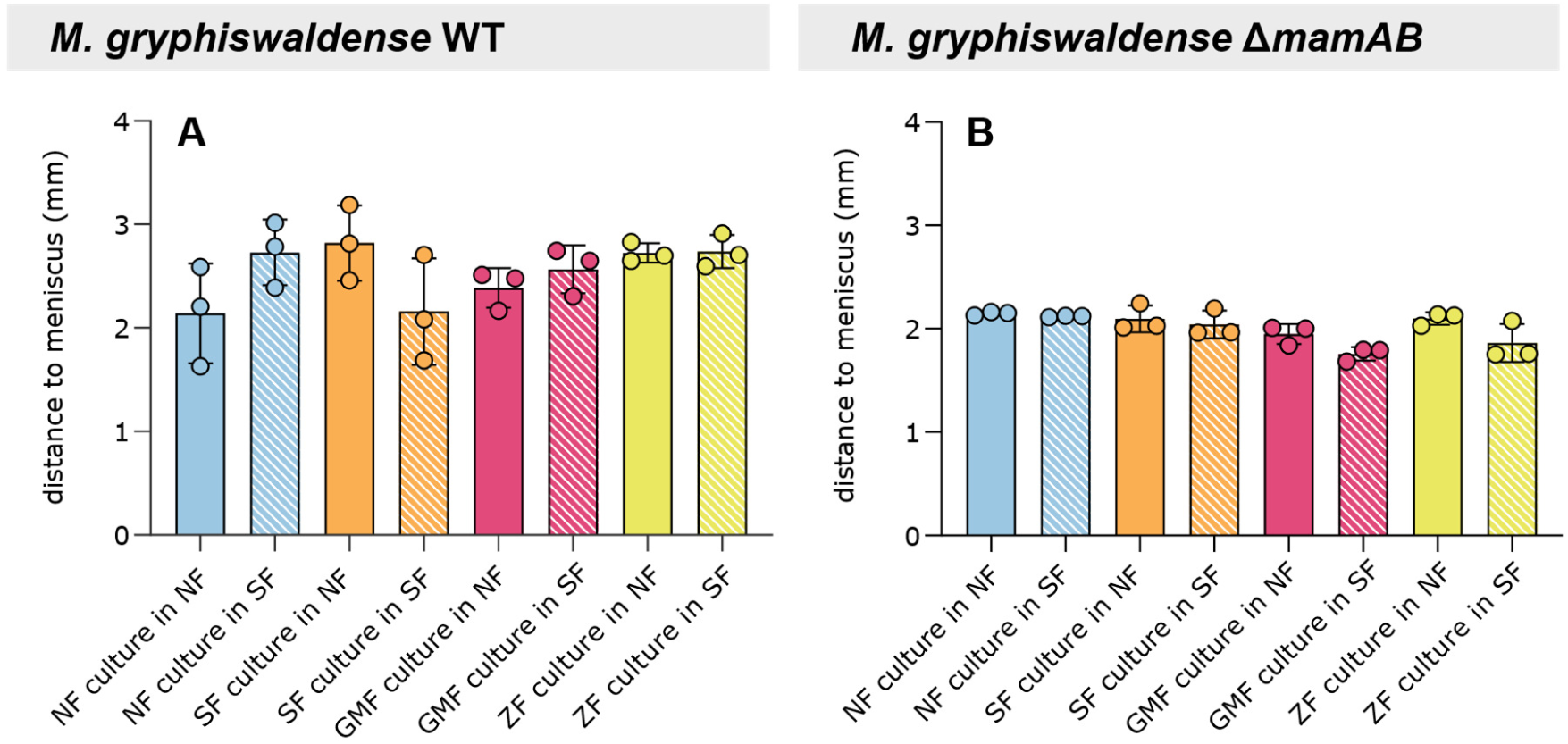
Distance of the aerotactic band to the air-liquid interface within the microcapillary after 180 min for *M. gryphiswaldense* wild-type (A) and Δ*mamAB*. **(B)** cultures. Overall, no statistically significant differences were found between samples and magnetic field conditions (Kruskal-Wallis test with Dunn’s multiple comparison test, *p* ≥ 0.05). However, aerotactic bands of NF cultures stabilized slightly closer to the meniscus under the NF setting, while SF-culture bands were similarly positioned under the SF setting **(A)**. These positional differences were not observable in Δ*mamAB* cultures **(B)**. Bars represent the mean and error bars the standard deviation (SD) from *n* = 3 independent experiments. Individual measurements are shown as dots.

To confirm that the differences observed among wild-type cultures in the capillary assay are due to differences in swimming polarity, we performed equivalent cultivation and capillary experiments using the non-magnetic Δ*mamAB* strain, which lacks the entire *mamAB* operon essential for magnetosome formation [21] and is therefore unable to develop swimming polarity (**Fig. S5**) [7]. Compared to wild-type cells (**Fig. S3**, **Fig. S4A**), Δ*mamAB* cells showed only minor differences in aerotactic band morphology and spatiotemporal dynamics during band formation under magnetic field influence (**Fig. S4B**, **Fig. S6**), indicating that aerotaxis alone is sufficiently efficient under the given experimental conditions. However, unlike the wild type, the Δ*mamAB* strain showed no differences in aerotactic band intensity distributions between cultures grown under different magnetic field configurations or between fields applied parallel versus antiparallel during the capillary assay (**Fig. 4**). This confirms that the differences observed among wild-type samples (**Fig. 2**) are attributable to the biomagnetism-dependent selection of swimming polarity during preceding culture growth.

**Fig. 4:**
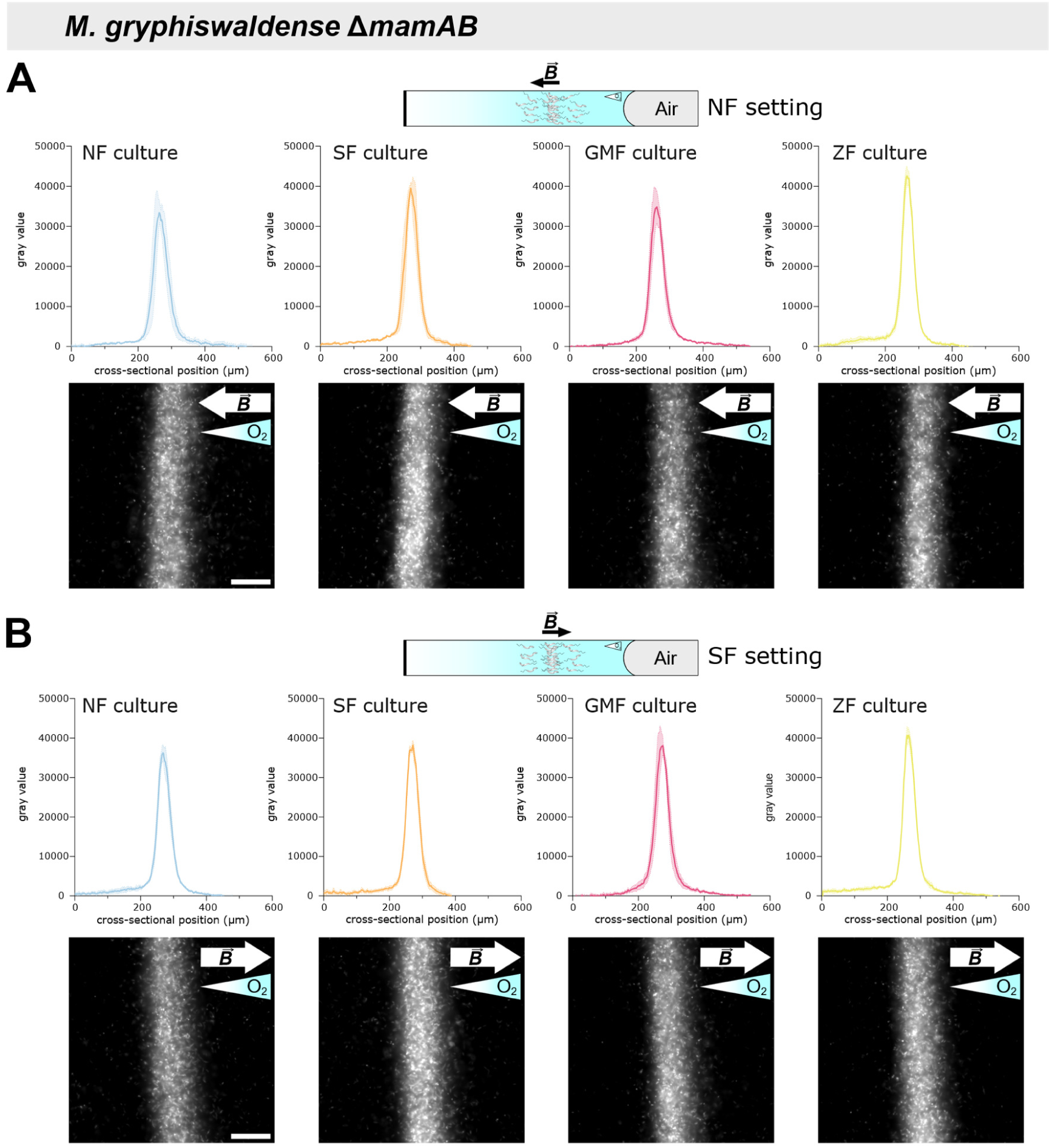
Aerotactic band intensity profile analysis of *M. gryphiswaldense* Δ*mamAB* cultures. Δ*mamAB* cells were pre-cultured and analyzed in microcapillaries using procedures identical to those applied to the wild-type strain. Band intensity profiles (mean ± SEM, *n* = 3) were determined from dark-field micrographs after 180 min. Unlike the wild type, Δ*mamAB* cultures showed consistent aerotactic band intensity profiles across all precultivation conditions. Arrows denote the direction of the magnetic field (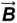), while gradient-filled triangles indicate oxygen gradient directions. Scale bars: 50 µm (apply to all images; shown only in the leftmost images).

### A Matching Polarity Bias Confers Competitive Advantage in Magneto-Aerotaxis

To directly assess the competitive advantage conferred by swimming polarity, we conducted competition experiments between fluorescently labeled north- and south-seeking cells. To this end, we introduced genes encoding soluble green (monomeric NeonGreen, mNG) and red (monomeric Cherry, mCherry) fluorescent proteins into both wild-type and Δ*mamAB* strains. Cultures of the resulting strains were then incubated again under conditions permissive for swimming polarity selection. In cultures of fluorescently labeled wild-type cells, this resulted in a notable enrichment of cells exhibiting the expected swimming polarity based on the applied magnetic field orientation during cultivation, as revealed by hanging drop assays (**Fig. S7**). In contrast, fluorescent Δ*mamAB* cells grown under conditions permissive for swimming polarity selection showed no preferential accumulation on either side of the drop (**Fig. S7**). Subsequently, green- and red-labeled wild-type cells grown under opposite polarity-selection conditions were mixed at a 1:1 ratio and examined again by the hanging drop assay, confirming that cells of each fluorescence predominantly migrated to opposite sides of the drop (**Fig. 5**).

**Fig. 5:**
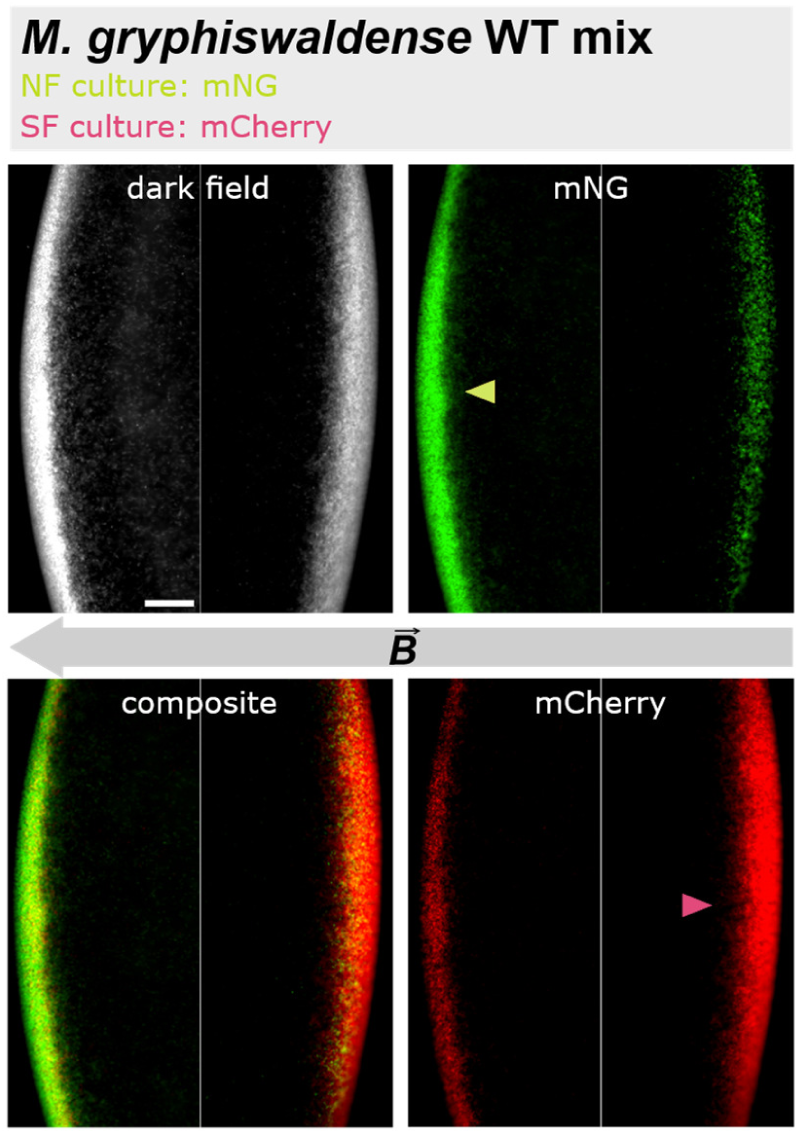
Swimming polarity-based separation of fluorescently labeled north- and south-seeking wild-type cells in the hanging drop assay. Predominantly north-seeking cells (NF culture), expressing mNG, and predominantly south-seeking cells (SF culture), expressing mCherry, were mixed in equal amounts. Green, north-seeking cells predominantly swam parallel to the applied magnetic field (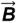), whereas red-labeled south-seeking cells primarily moved in the opposite, antiparallel direction. Scale bar: 50 µm (applies to all images; shown in the upper left image).

Mixed populations—comprising approximately equal numbers of green-labeled north-seeking and red-labeled south-seeking cells—were then subjected to capillary assays under NF and SF settings (**Fig. 6**). Aerotactic bands were imaged using sequential exposure with both fluorescence channels to quantify the abundance of green and red cells. When the NF setting was applied during the assay, green-fluorescent wild-type cells predominated within the aerotactic band, as evidenced by a stronger mNG signal compared to mCherry (**Fig. 6A**). In contrast, when the same mixed culture was analyzed under the SF settings, the opposite pattern emerged, and red-fluorescent wild-type cells accumulated at higher intensities within the band (**Fig. 6B**). These findings are consistent with capillary assays performed on non-fluorescent cells and confirm that correct swimming polarity confers a direct competitive advantage under matching magnetic field conditions. Consistent with this, no such effect was observed in corresponding assays with fluorescently labeled, non-magnetic Δ*mamAB* cells lacking a defined swimming polarity (**Fig. S8**). In these experiments, red and green fluorescence intensities—and thus cell abundances—were comparable within the aerotactic band, regardless of the applied magnetic field orientation (**Fig. 6C,D**).

**Fig. 6:**
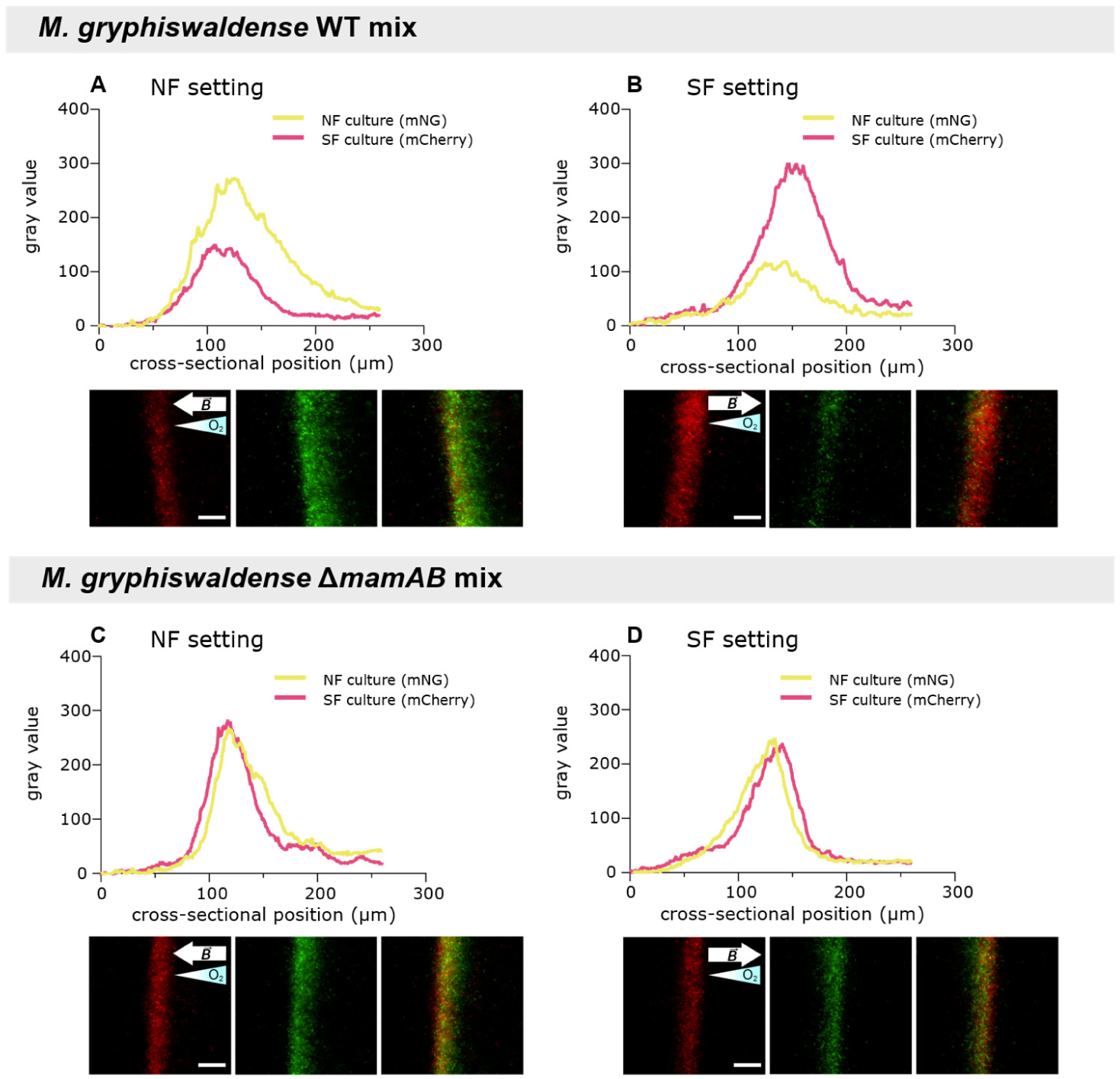
Swimming polarity confers a competitive advantage in mixed populations of north- and south-seeking cells. Equal proportions of green-fluorescent (mNG-expressing, predominantly north-seeking) and red-fluorescent (mCherry-expressing, predominantly south-seeking) wild-type cells were mixed and subjected to capillary assays under NF and SF settings. After 120 min, under the NF setting, green-fluorescent wild-type cells predominated in the aerotactic band **(A)**, whereas under the SF setting, red fluorescent cells outcompeted green-fluorescent cells **(B)**. In contrast the abundance of red and green Δ*mamAB* cells did not differ with respect to the applied magnetic field **(C, D)**. Shown are representative intensity profiles and fluorescence micrographs of aerotactic bands obtained from a total of *n* = 6 experiments. Arrows indicate the direction of the magnetic field (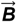), while gradient-filled triangles indicate the direction of oxygen gradients. Scale bars: 50 µm (apply to all images; shown only in the leftmost images).

### M. gryphiswaldense Exhibits Light-Driven Phototactic Responses

Several studies suggested that phototactic responses play an important role in MTB navigation, complementing magneto-aerotaxis [5, 22–24]. However, to our knowledge, until now, phototactic behavior in *M. gryphiswaldense* had not been reported. Experiments with fluorescence illumination revealed an interesting side finding: the aerotactic band shifted by more than 100 µm toward the meniscus upon activation of the fluorescence light source using a green fluorescent protein (GFP) filter set (i.e., blue light). Migration of the band toward the meniscus occurred rapidly, irrespective of whether the NF or SF setting was applied during the experiment, with a change in position within a few seconds, after which movement slowed. This behavior was observed not only in wild-type populations (**Fig. 7**, **Movie S1**) but also in non-magnetic Δ*mamAB* cells (**Movie S2**), further indicating that it is independent of biomagnetism. Notably, the blue-light response was only partially reversible—when the fluorescence illumination was switched off, the aerotactic band returned within a few seconds to a position closer to, but not identical with its original location (**Fig. 7**, **Movies S1–2**). No band shift was observed when green light (mCherry filter set) was switched on; however, subsequent switching off the green light caused a slight shift of the band away from the meniscus (**Movie S3**). During continuous blue-light exposure (∼12 min), the aerotactic band steadily migrated toward the meniscus, but its movement slowed over time and stabilized at ∼5 min (**Movie S4**), indicating cellular adaptation to constant blue-light stimulation. It should be noted, however, that after prolonged blue light exposure, the band also partially dissipated, suggesting potential harmful effects on the cells that prevented further migration of the band.

**Fig. 7:**
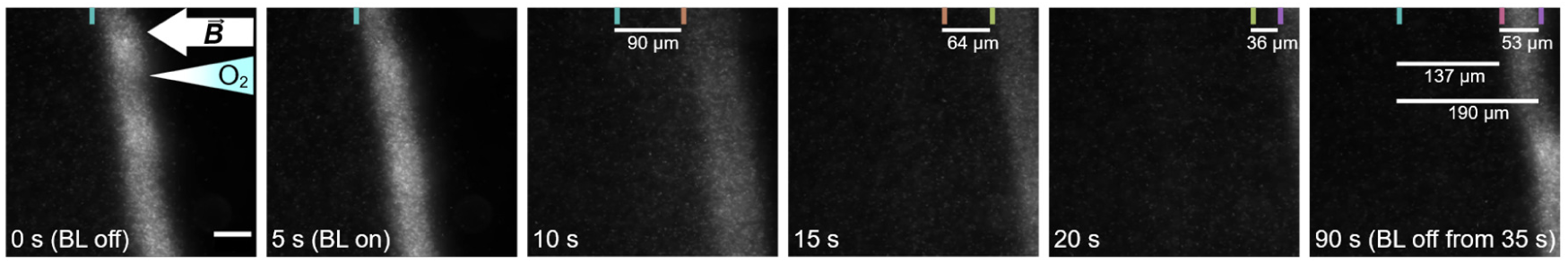
Phototactic response to blue light in *M. gryphiswaldense.* Individual frames from a representative time-lapse recording of wild-type cells using dark-field illumination are shown. The excitation light source for fluorescence imaging (GFP filter set) was activated at 5 s and switched off at 35 s. Blue light (BL) induced a unidirectional movement of cells toward the meniscus, suggesting that phototactic behavior is mechanistically coordinated with aerotaxis. Colored bars indicate the position of the aerotactic band, with each color assigned to a specific location. When the band shifts between time points, the bar color changes accordingly. The movement of the band between two images is visualized by the spatial separation of bars of different colors in the respective frames. Time stamps are indicated in each image. The white arrow shows the direction of the magnetic field (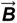), while the colored triangle indicates oxygen gradient direction. Scale bar, 50 µm, in the leftmost image applies to all images.

## Discussion

The ability of organisms to respond to Earth’s magnetic field, known as magnetosensation, was likely shaped by evolutionary pressures arising from the dynamic geomagnetic field, which varies in intensity and undergoes complete polarity reversals over geological time scales [4, 14, 25, 26]. MTB not only align with magnetic fields of a given strength but also respond to field polarity—a behavior closely linked to their chemosensory abilities that allow for temporal perception of oxygen gradients [4, 7–9, 20]. Here, we examined the impact of this polar response on magneto-aerotaxis. We show that matching swimming polarity is crucial for directing cells toward low-oxygen environments, as evidenced by their predominant accumulation within the aerotactic band only when the magnetic field orientation relative to the oxygen gradient in the capillary matches that experienced during prior culture incubation (**Fig. 2**, **Fig. 6**). Together with magnetic field inversion experiments [5, 8], these findings suggest that cells with a non-matching polarity bias are redirected into regions above or below their microaerophilic optimum. Unlike the oxygen gradients observed in microcapillaries [9], the hanging drop assay—when performed with sufficiently small droplets exposed to air—creates an environment where oxygen levels likely exceed the physiological optimum due to rapid diffusion from the atmosphere [6, 17]. This is ultimately expected to trigger a repellent response, causing cells to accumulate at the droplet edges—preferentially on the side facing magnetic north or south, depending on their polarity bias. Our results further show that combining microcapillary assays with intensity profile analysis allows quantification of north- and south-seeking subpopulations (**Fig. 2**). In contrast, the hanging drop assay is, at best, semi-quantitative, as pixel intensity measurements at the drop edge are prone to error due to the curvature of the droplet, making results sensitive to the focal plane. Whereas the peak height (or, alternatively, the area) of the aerotactic band intensity profile is well suited for quantifying swimming polarity (**Fig. 2**), the band width (**Fig. S4**) and the distance to the meniscus appear less suitable (**Fig. 3**). In particular, the latter is influenced by the steepness of the oxygen gradient, which in turn depends on the number of cells present in the band and their respiratory activity [9, 18]. The resulting altered spatio-temporal dynamics of band formation may explain the—rather small and statistically non-significant—differences in band distance to the meniscus between wild-type populations with non-aligned swimming polarity, those with aligned swimming polarity, and Δ*mamAB* cells (**Fig. 3**).

Despite general agreement with previous studies, some unclarities remain. For instance, our analysis suggests that zero-field-grown *M. gryphiswaldense* populations contain equal fractions of north- and south-seeking cells (**Fig. 1**, **Fig. 2**). This implies that, in the presence of a magnetic field, half of the cells—those with non-matching polarity—are guided away from the aerotactic band and do not contribute to its formation, as reflected by reduced peak intensities in aerotactic band profiles (**Fig. 2**). Consistent with our observations, early studies reported equal numbers of MTB of each swimming polarity in mud samples maintained in a zero-field environment for extended periods [1, 12]. Moreover, this distribution is consistent with what has been observed in hanging drop assays of (GMF-exposed) *M. gryphiswaldense* cultures grown under homogenous oxygen concentration [7]. However, field inversion experiments on *M. gryphiswaldense* populations grown under homogeneous oxygen concentrations—as well as similar experiments with “axial” *Magnetospirillum magnetotacticum* populations, which form aerotactic bands at both ends of a capillary opened at both ends (unlike polar magnetotactic cocci that formed bands only at one end of the capillary)—showed that cells rotated by 180° upon field inversion but remained within the band [5, 6, 8, 17]. Thus, different results provide conflicting information about whether a genuine “axial” state exists, or if “axial” merely refers to populations containing equal numbers of north- and south-seeking cells. Since our study design did not include field inversion experiments, we cannot comment on whether the polarity bias of cells among different cultivation samples after band establishment differed from their initial bias prior to band formation. Furthermore, comparing results across studies is challenging due to variations in growth conditions and sample preparation. Nevertheless, the previous simultaneous observation of *M. gryphiswaldense* cells swimming away from the band in both directions, alongside cells that rotate by 180° and remain within the band upon field reversal, indicates the coexistence of subpopulations of “polar” and truly “axial” cells [8]. In addition, although hanging drop assays indicated an equal proportion of north- and south-seeking cells, the observed behavioral responses to oxygen upshift in non-polarized *M. gryphiswaldense* populations further imply they are not simply a mixture of both swimming polarities [7].

Another important point to consider is that, although previous capillary experiments with wild-type cells under zero-field conditions and magnetic fields (anti)parallel to the oxygen gradient indicated a general benefit of magnetic alignment for aerotactic band formation [9, 18, 27], we did not observe pronounced differences between wild-type and non-magnetic Δ*mamAB* cells in the capillary assay (**Figs. 2–4**, **Figs. S2–4**, **Fig. S6**). These findings suggest that aerotaxis alone is sufficiently effective under the given experimental conditions. Notably, the navigational benefit of magnetic alignment may be greater in structured environments that resemble natural MTB habitats (where movement is spatially constrained) [28, 29] compared to bulk liquid conditions in capillary assays, in systems subject to perturbations [5], or in long-term cultivation experiments [30]. Moreover, it must be considered that even within our polarity-selected wild-type populations, a certain fraction of cells consistently exhibited non-matching polarity (**Figs. 1–2**). Therefore, to draw definitive conclusions, similar experiments—including direct competition assays—should be conducted using not just comparable overall numbers of wild-type and Δ*mamAB* cells, but specifically wild-type cells of a single polarity type. Achieving such populations through a cultivation-dependent approach is experimentally challenging (**Figs. 1–2**), but could potentially be accomplished via magnetic enrichment of cells with one polarity type following selective growth [8].

Additionally, our results provide evidence that a stronger, artificial magnetic field imposes a more pronounced selective pressure on swimming polarity (**Fig. 1**). Nonetheless, consistent with observations that fresh environmental magnetospirilla isolates display swimming polarity [6, 31–33], we experimentally confirm here that the GMF is sufficient to drive north-seeking polarity selection in *M. gryphiswaldense*. Given that redox gradients in natural sediments are more stable than in our GMF-exposed laboratory cultures, the selective pressure in natural environments is likely stronger. In accordance with prior work [7, 34], we also observed that swimming polarity in *M. gryphiswaldense* lab cultures is selected within relatively few generations (**Fig. 1**), although even extended cultivation did not yield populations consisting exclusively of a single polarity type. Compared to the rapid adaptation of swimming polarity in *M. gryphiswaldense*, in cultures of other MTB it took several weeks of incubation for the inversion of the population-wide swimming polarity bias to occur [1, 6, 12, 17]; however, this process is still extremely rapid when considered on geological time scales. Given that geomagnetic field reversals take thousands of years to complete, these findings raise questions about the selective mechanisms driving such rapid adaptive behavior [25, 26].

Currently, the molecular mechanisms governing swimming polarity determination and inheritance in single cells, as well as the selection of the prevailing polarity type at the population level, are poorly understood. In *M. gryphiswaldense*, it is known that swimming polarity is closely tied to aerotaxis and the chemosensory pathway encoded by the major chemotaxis operon 1 [7]. Moreover, swimming polarity may depend on motility-related cellular asymmetries (cellular polarity)—such as a monopolar flagellum (see, e.g., Figs. 1 and 3 in ref. [6]) or two structurally distinct counterrotating flagellar motors in bipolarly flagellated magnetospirilla (**Fig. 8**)—and on how these structures align with the overall magnetic dipole (magnetic polarity) generated by the combined magnetosome dipoles. In cells where cellular polarity is inherited to daughter cells during division (**Fig. 9i**), inversion of swimming polarity might be initiated by random loss of magnetosomes and inversion of magnetic polarity by *de novo* formation of magnetosomes (**Fig. 9ii**), with a 50% probability of magnetic polarity inversion upon magnetosome loss [12, 17, 25]. However, magnetosome chain distribution to daughter cells is tightly regulated in *M. gryphiswaldense* [35, 36]. Apart from a loss of magnetosomes, remagnetization and swimming polarity inversion can be triggered by strong magnetic pulses (in the range of several tens of millitesla) [1, 37]. Although this may be relevant when permanent magnets are used to investigate or enrich MTB [38], such field intensities are unlikely in natural environments. Alternatively, swimming polarity inversion might occur as a result of the production of heterogeneous offspring as a bet-hedging strategy (**Fig. 9iii**). It has been suggested that remaining cells of the opposite swimming polarity (as also observed in our experiments, **Fig. 1**) might serve as progenitors and enable rapid polarity inversion at the population level upon a field reversal [1]. Subsequently, the dominant swimming polarity type could be selected by the prevailing orientation of the magnetic field relative to the oxygen gradient. However, this strategy may be energetically costly, as a fraction of the population would be consistently directed into hypo- or hyperoxic regions unfavorable for growth. The simultaneous occurrence of MTB with opposing swimming polarities in environmental samples and laboratory cultures [1, 8, 10, 15, 16] could also be an indicator that swimming polarity selection is influenced by multiple environmental cues, including redox and oxygen gradients (both downward and inverted), light (see below), or geomagnetic anomalies from magnetic grains or rocks [1, 5, 6, 8, 27, 39–41]. As discussed above, swimming polarity may also arise gradually within individual cells from an intermediate non-polar (or genuine axial) state (**Fig. 9iv**), either in a selective manner or controlled by a dedicated, switchable molecular mechanism that regulates cellular polarity [7]. Potential mechanisms underlying swimming polarity determination are currently under investigation in our laboratory.

**Fig. 8:**
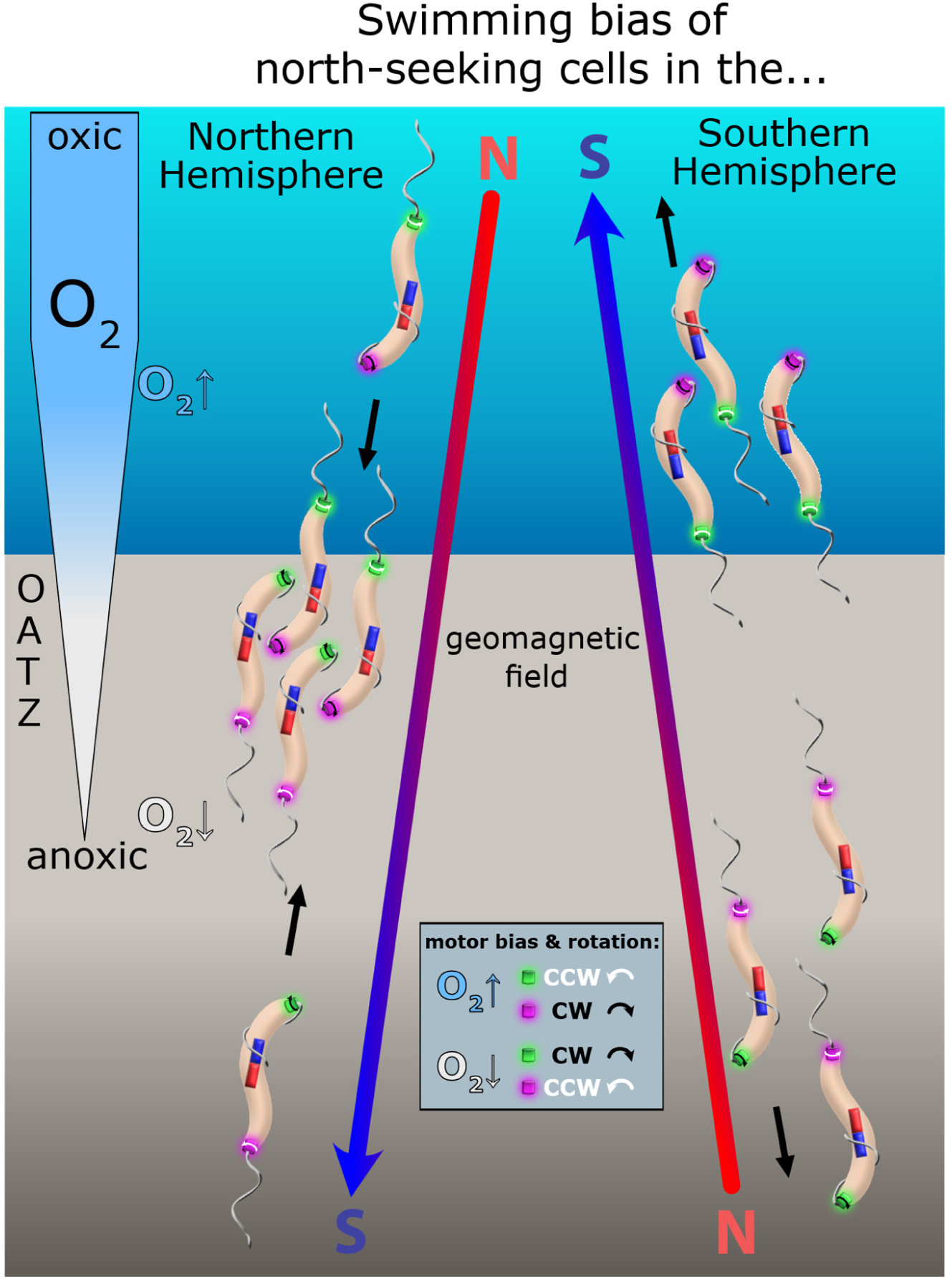
Hypothetical model of swimming polarity determination in bipolarly flagellated magnetospirilla. In the illustrated model, swimming polarity is governed by the spatial positioning of two structurally distinct, counter-rotating flagellar motor subtypes relative to the net magnetic moment of the magnetosome chain (represented by bar magnets). The two motors (shown in green and purple) exhibit opposing switching behavior when cells encounter oxygen concentrations deviating from their microaerophilic optimum. Exposure to oxygen levels exceeding the optimum (O2↑) increases the probability of switching—the purple motor tends to switch to clockwise (CW) rotation, while the green motor switches to counterclockwise (CCW) rotation. Under suboxic conditions (O2↓), the opposite switching behavior occurs. CW motor rotation is consistently associated with a wrapped flagellum at the leading cell pole, whereas CCW motor rotation corresponds to a lagging pushing flagellum. As a result, north-seeking polarity cells (only these are depicted) swim toward the oxic–anoxic transition zone (OATZ) under Northern Hemisphere magnetic field conditions, but away from the OATZ under Southern Hemisphere conditions. The model defines swimming polarity solely by the orientation of magnetic polarity relative to cellular polarity. North- and south-seeking cells respond similarly to temporal changes in oxygen levels, and the model does not incorporate magnetoreceptive mechanisms that involve sensing magnetic field intensity or polarity. Note that this model is partly based on existing literature but also incorporates experimentally unverified assumptions. It is primarily intended to illustrate how our findings in this study may be explained at the level of individual cells.

**Fig. 9:**
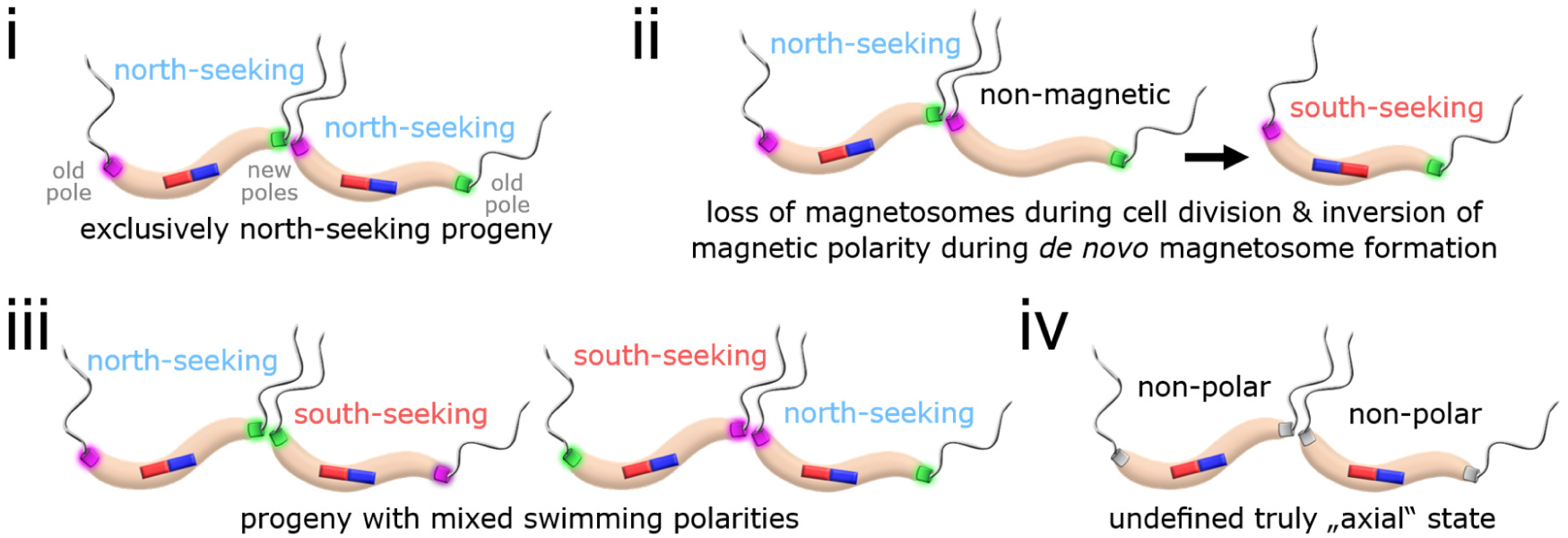
Hypothetical mechanisms for swimming polarity inheritance and inversion in bipolarly flagellated magnetospirilla. **(i)** Swimming polarity may be maintained through the inheritance of fixed positioning of two flagellar motor subtypes relative to the net magnetic moment of the magnetosome chain. Note that, relative to the old and new cell poles (indicated in gray font), daughter cells exhibit opposing cellular polarities to account for their opposing magnetic polarities when magnetosome chains are split during cell division. **(ii)** If swimming polarity is maintained through cell division, swimming polarity inversion may occur due to random loss of magnetosomes during division and subsequent *de novo* magnetosome formation (i.e., inversion of magnetic polarity). **(iii)** Alternatively, each cell division may generate offspring comprising both swimming polarity types. **(iv)** Swimming polarity might also develop from an undefined truly axial state. Note that north- and south-seeking swimming polarity here refers not only to the immediate observed swimming direction of cells, as in some studies, which depends on whether cells encounter oxygen levels above or below their optimum, but to the overarching swimming polarity type characteristic of the cells. This figure incorporates unverified assumptions and is intended to illustrate potential mechanisms of swimming polarity inversion.

Notably, we also discovered a biomagnetism-independent, partially reversible, phototactic response in *M. gryphiswaldense*, characterized by the unidirectional collective movement of cells toward higher oxygen concentrations upon blue-light exposure (**Fig. 7**, **Movies S1–2**, **Movie S4**). The collective upward movement of *M. gryphiswaldense* cells in an oxygen gradient indicates that the light-dependent response is linked to aerotactic behavior. Similar observations were made in capillary experiments with magnetococcus strain MC-1, but not when using *Magnetospirillum magnetotacticum* [5]. However, in this study, a migration of polar MC-1 cells away from the meniscus was observed upon blue-light exposure [5]. A blue-light response was also recently identified in the *M. gryphiswaldense*-related strain *Paramagnetospirillum magneticum* AMB-1 (via U-turn analysis) and attributed to the LOV-domain protein Amb2291 and the methyl-accepting chemotaxis protein Amb0994 [24]. Since the *M. gryphiswaldense* genome encodes similar proteins [42, 34], it will be important in the future to analyze their role in phototaxis. It will also be crucial to understand the physiological relevance of this response in natural habitats and how it integrates with aerotaxis. Given the large penetration depth of blue light in water, it remains unclear why cells would swim toward higher oxygen levels upon blue light exposure—potentially exposing themselves to even higher doses of blue light. Also, the band shift observed after cessation of green light exposure (**Movie S3**) may indicate additional mechanisms underlying light-driven spatial environmental orientation, warranting further investigation.

## Conclusions

Our study underscores the importance of the geomagnetic field in magnetic-pole-dependent directional motility and highlights the adaptive nature of polar magneto-aerotaxis. We demonstrate that this behavioral response is tightly coupled with aerotaxis, offering a quantitative measure of its role in navigating toward low-oxygen environments. Moreover, we show that *M. gryphiswaldense* integrates light-dependent stimuli into its aerotactic responses. Collectively, these findings advance our understanding of bacterial motility and emphasize the sophisticated sensing and integration of environmental cues in *M. gryphiswaldense*. They may also inform bioengineering strategies that exploit MTB as microrobots guided by oxygen, magnetic fields, and light-based (opto) cues, enabling precise spatiotemporal control of their movement for applications in microscale transport, targeted delivery, or environmental sensing.

## Methods

### Cultivation of Bacterial Strains

*M. gryphiswaldense* was cultivated in modified flask standard medium (FSM) [43] at 28°C. Selection of swimming polarity was carried out according to established protocols [7]. This involved exposing non-agitated cultures (over subsequent culture passages) in glass screw-cap tubes with caps loosened by half a turn for at least 10 and up to 30 generations, to either a uniform 0.6 mT magnetic field applied parallel or antiparallel to the oxygen gradient (simulating Northern or Southern Hemisphere geomagnetic field polarity, respectively) using coils, the local geomagnetic field (Bayreuth, Germany with north, east, and vertical components of 20.0 µT, 1.5 µT, and 45.0 µT, respectively), or a zero-field generated by triaxial pairs of coils as a control (**Fig. S1A**). The optical density and magnetic response (*C*_mag_) of *M. gryphiswaldense* cultures were measured at 565 nm as described previously [34]. *Escherichia coli* was grown in lysogeny broth (LB) at 37°C with shaking at 180 rpm. For the cultivation of *E. coli* WM3064 (W. Metcalf, unpublished), 0.1 mM DL-α,Ɛ-diaminopimelic acid (DAP) was added. Media were solidified by adding 1.5% (w/v) agar. Selective growth was achieved using kanamycin at a concentration of 5 µg/ml for *M. gryphiswaldense* and 25 µg/ml for *E. coli*. All strains are listed in **Table S1**.

### Molecular and Genetic Techniques

Oligonucleotides (listed in **Table S2**) were obtained from Sigma-Aldrich (Steinheim, Germany). Genes of interest were amplified using Q5 (NEB) proofreading DNA polymerase. Plasmids (**Table S1**) were constructed using standard molecular biology techniques, incorporating FastDigest™ restriction enzymes and T4 DNA Ligase (Thermo Scientific). Constructs were sequenced by Macrogen Europe (Amsterdam, Netherlands). For the construction of mNG- and mCherry-producing *M. gryphiswaldense* strains, a *M. gryphiswaldense* codon-optimized *mNeonGreen* gene (*omNG100*) and the *mCherry* gene, each under the control of the constitutive P*_mamDC45_* promoter, were amplified using primer pairs 717/718 and 717/719, respectively, and cloned into the Tn*7* vector pBAMII, which enables site-specific genomic insertion [34]. The resulting constructs were transferred from *E. coli* WM3064 into the *M. gryphiswaldense* wild-type and Δ*mamAB* strains via conjugation. Successful transconjugants were verified by PCR screening using primers 558/559.

### Hanging Drop Assay

Swimming polarity of *M. gryphiswaldense* populations was assessed using a modified version of the classical hanging-drop assay. Following cultivation, microdroplets (0.8 µL) of cell suspension were transferred onto a coverslip, which was placed upside down on a stack of five layers of laboratory tape and imaged by dark-field illumination and/or fluorescence using the same microscope [20] used for the microcapillary assays (see below). To assess swimming polarity, a homogeneous 400 µT horizontal magnetic field was applied using electromagnetic coils [20], and images of the droplet edges facing magnetic north and south were taken after ∼5 min.

### Microcapillary Assays

Analysis of aerotactic band formation under defined magnetic field conditions was performed using custom-made triaxial magnetic coils mounted on a Nikon Eclipse FN1 upright microscope at room temperature. The microscope was equipped with a Nikon S Plan Fluor ELWD 20× DIC N1 objective (NA 0.45), a Nikon dark-field condenser (NA dry 0.80 - 0.95), and a pco.edge 4.2 sCMOS camera (PCO). For the experiments, culture samples were adjusted to an OD_565_ of 0.1, and 1 ml of the diluted suspension was purged with N_2_ for 15 min. Samples were transferred into rectangular glass capillaries (0.1 × 1 mm; Vitro Tubes #5010-050) by capillary forces. Capillaries were sealed at one end with vacuum grease. To study cell behavior under homogeneous 400 μT magnetic fields mimicking Northern- and Southern-hemisphere geomagnetic field polarity relative to the oxygen gradient, two capillaries were filled with the same cell suspension and positioned adjacent to each other, with one capillary inverted by 180° so that the aerotactic bands were at approximately similar positions along the microscope stage x-axis (**Fig. S1B**). Images and/or videos were captured at various time points to document the spatiotemporal dynamics of aerotactic band formation. Images were acquired in dark-field illumination with a camera exposure of 40 ms, a readout speed of 95 MHz, and the manual light source intensity set to 11 o’clock to prevent overexposure. Band distances from the meniscus and intensity profiles were measured using the line length and profile tools in ImageJ Fiji [44]. For competition experiments, equal numbers of fluorescently labeled wild-type and Δ*mamAB* cells were mixed, and the ∼1:1 ratio was verified by fluorescence microscopy. Capillaries were filled with mixtures, and the capillary assay was performed as described above. Fluorescence illumination and acquisition were performed using a SOLA SE II light source (Lumencor) set to 100% intensity with 200 ms exposure time. The following filter sets were used: GFP (excitation 482/18 nm, dichroic 495 nm, emission 520/28 nm) and mCherry (excitation 562/40 nm, dichroic 593 nm, emission 640/75 nm). To minimize both the blue-light-dependent aerotactic band shift and photobleaching, images of the aerotactic band in fluorescence mode were acquired immediately after activation of the excitation illumination.

### Statistical Analysis

Statistical analyses were conducted using GraphPad Prism 10.1.1. Details on sample sizes (*n*), statistical tests, and *p*-value definitions are provided in the figure legends.

## Supporting information

Legends for Supplementary Movies

## List of abbreviations

MTB: magnetotactic bacteria
NF: northern field
SF: southern field
GMF: geomagnetic field
ZF: zero field
WT: wild type
mNG: monomeric NeonGreen
mCherry: monomeric Cherry
GFP: green fluorescent protein
BL: blue light
SEM: standard error of the mean
SD: standard deviation

## Declarations

### Ethics approval and consent to participate

Not applicable

### Consent for publication

Not applicable

### Availability of data and materials

The datasets used and/or analyzed during the current study are available from the corresponding author on reasonable request.

### Funding Acknowledgment

This work was funded by the Deutsche Forschungsgemeinschaft (DFG, German Research Foundation)—Project number 525457187, and Project number 521548282 under Germany’s Priority Program “Reconstructing the deep dynamics of planet Earth over geologic time (DeepDyn)” SPP2404 – 500707704.

### Authors’ contributions

D.P. designed the study. C.W. performed experiments. C.W. and D.P. analyzed data. C.W. and D.P. wrote the manuscript.

