## Supplementary Figures for "Directional Matching of Swimming Polarity Provides a Competitive Advantage During Bacterial Magneto-Aerotaxis"

**Additional materials for**

**This PDF file includes:**

Figures S1 to S8

**Other additonal materials for this manuscript include the following:**

Tables S1 to S2
Legendes for Movies S1 to S4
Movies S1 to S4

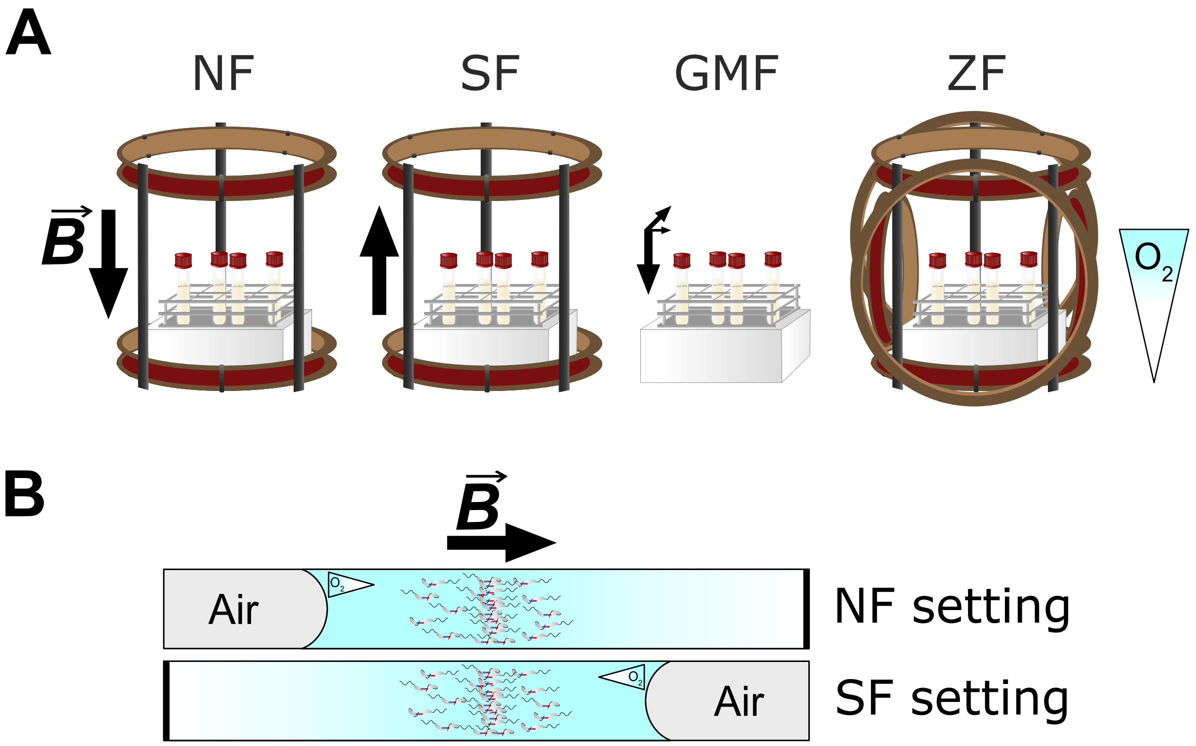


**Fig. S1. Magnetic field configurations used during culture incubation and in capillary assays. (A)** Preceding cultures were grown under four different magnetic field conditions in non-agitated culture tubes, allowing an oxygen gradient to form: NF: Northern Field, a uniform 0.6 mT field mimicking Northern Hemisphere polarity (generated by coils). SF: Southern Field, a uniform 0.6 mT field mimicking Southern Hemisphere polarity (generated by coils). GMF: The local geomagnetic field in Bayreuth, Germany (north, east, and vertical components of 20.0 μT, 1.5 μT, and 45.0 μT, respectively; horizontal field ≈ 20.1 μT; total magnitude ≈ 49.3 μT). ZF: A zero field produced by triaxial coil pairs. **(B)** To compare cell behavior under magnetic fields resembling Northern- and Southern-Hemisphere geomagnetic field polarity (referred to as NF and SF setting, respectively), two capillaries filled with cell suspensions were placed side-by-side, with one rotated by 180° to align the aerotactic bands along the microscope stage x-axis under a homogeneous 400 μT magnetic field. Black arrows indicate the predominant field direction and polarity, and gradient-filled triangles show the direction of the oxygen gradients.

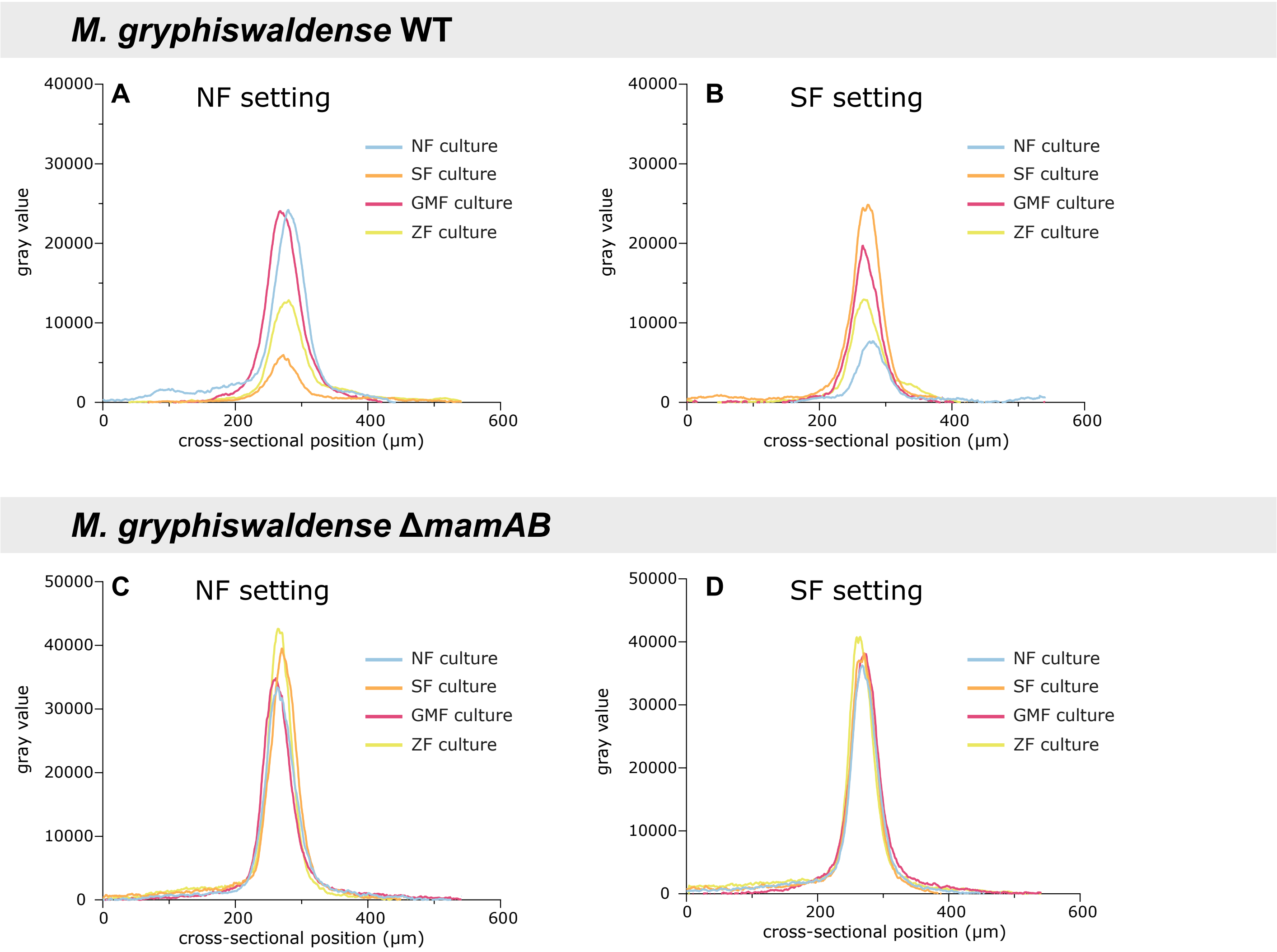


**Fig. S2. Aerotactic band intensity profiles.** Overlaid, averaged aerotactic band intensity profiles (after 180 min) from NF, SF, GMF, and ZF cultures of both the wild type **(A, B)** and the Δ*mamAB* strain **(C, D)** under NF **(A, C)** and SF **(B, D)** settings. Data correspond to **Figs. 2** and **4**.

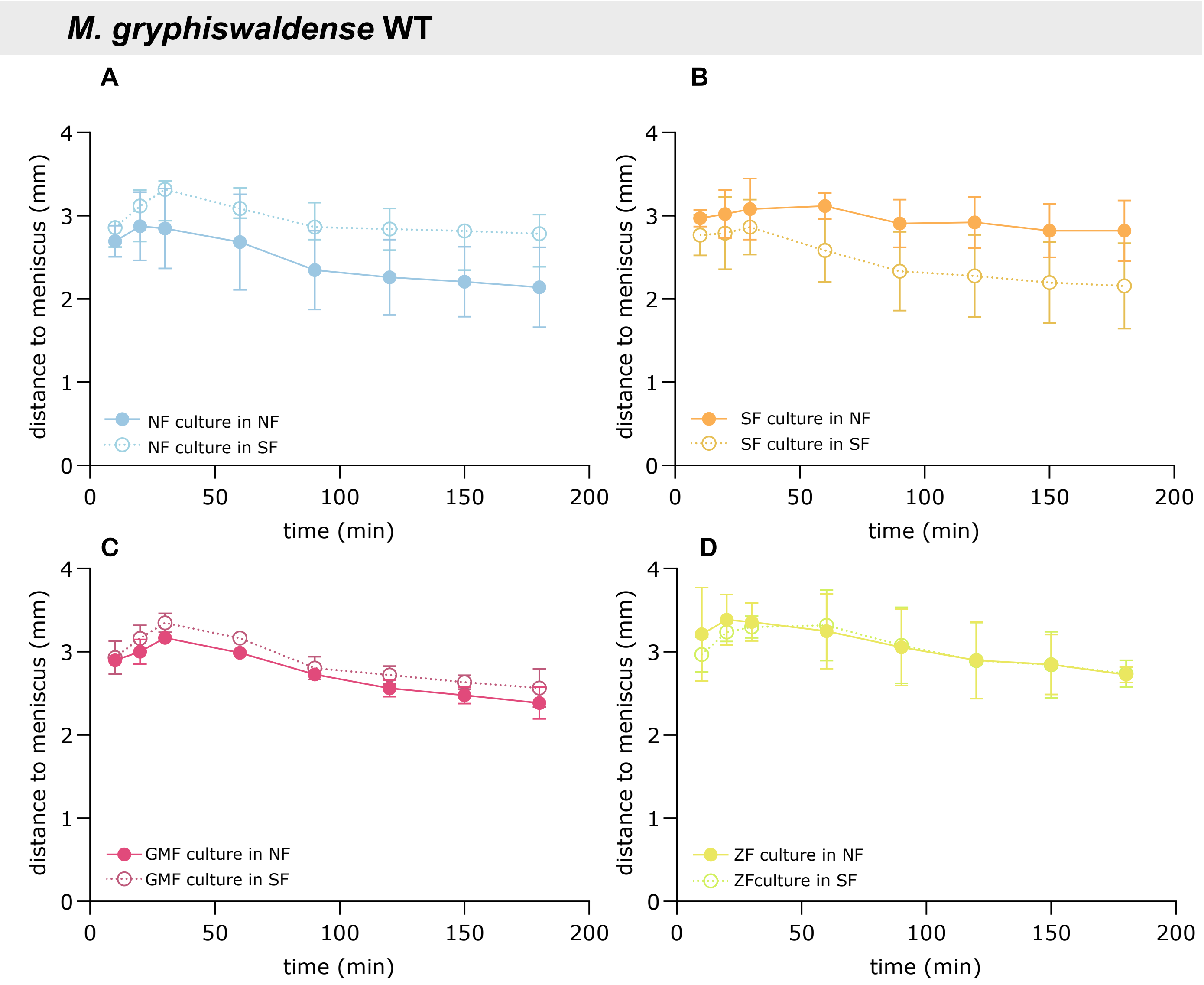


**Fig. S3. Spatiotemporal dynamics of aerotactic band formation for wild-type cultures.** The mean distance (± SD) between the aerotactic band and the air-liquid interface (meniscus) was measured at defined time points (*n* = 3 capillary experiments). **(A)** NF culture; **(B)** SF culture; **(C)** GMF culture; **(D)** ZF culture—each analyzed under NF and SF settings (solid and dashed lines, respectively). Data correspond to **Figs. 2** and **3**.

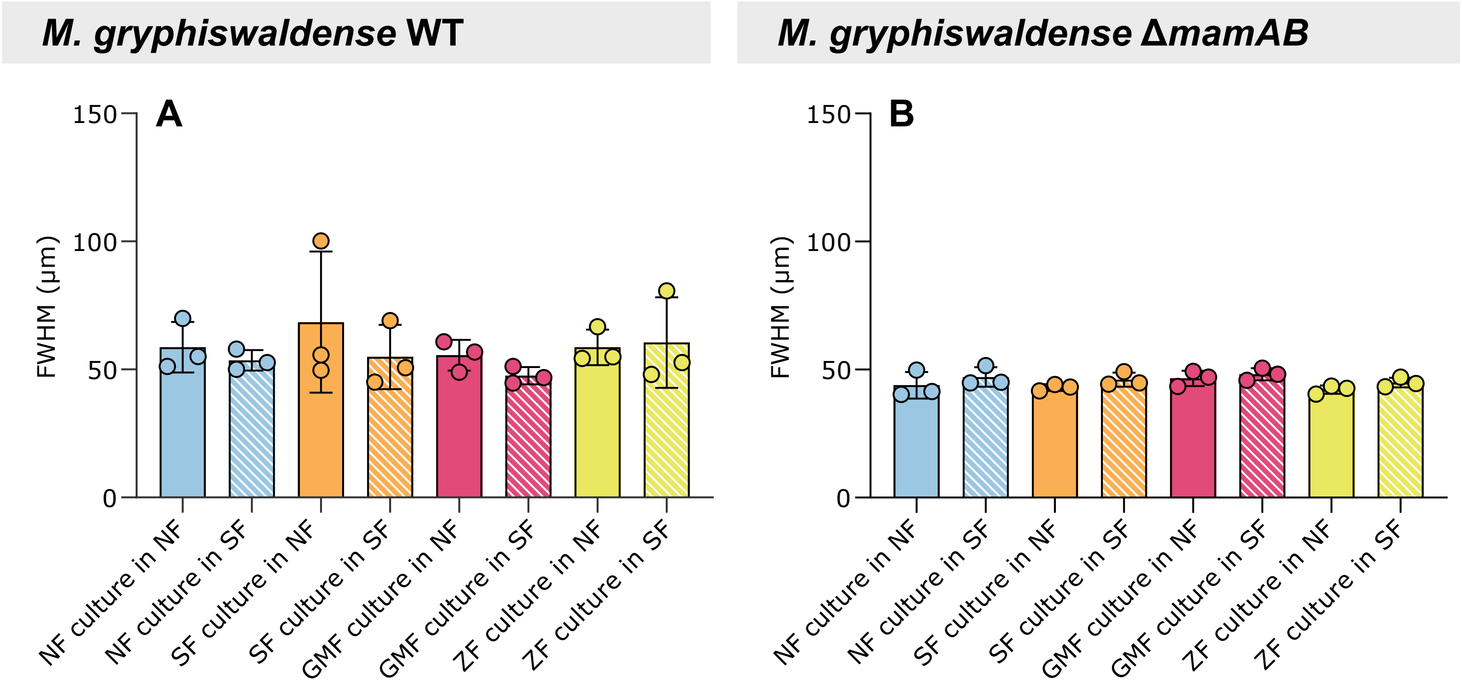


**Fig. S4. Widths of aerotactic bands.** The full width at half maximum (FWHM) of aerotactic bands was measured after 180 min for the wild-type **(A)** and Δ*mamAB* strain **(B)** under both NF and SF settings (colored and dashed bars, respectively). Bars represent the mean, error bars the SD, and individual measurements are shown as dots ($n$ = 3 independent experiments). No statistically significant differences were found between samples or magnetic field conditions (Kruskal-Wallis test with Dunn's multiple-comparison test, $p \geq 0.05$). Data correspond to **Figs. 2–4**.

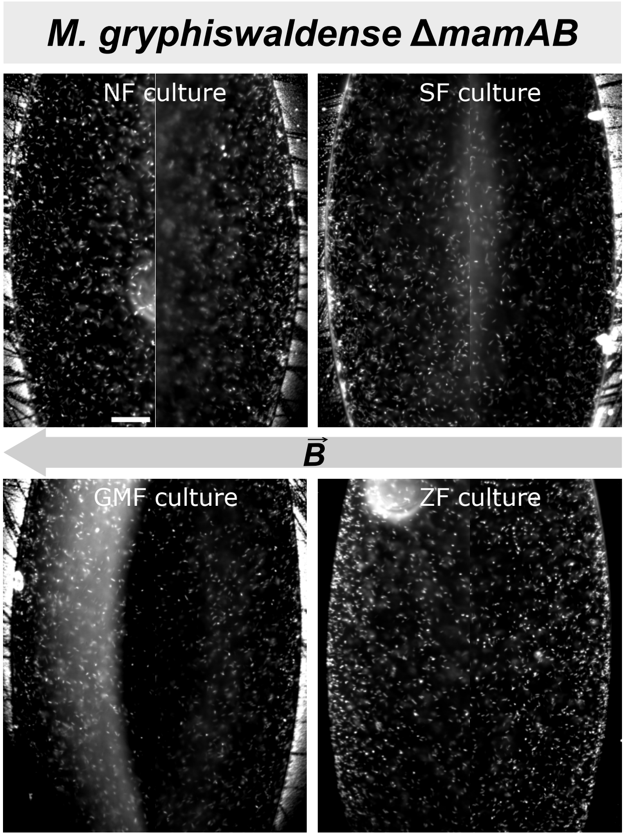


**Fig. S5. Hanging-drop assay of Δ*mamAB* cultures.** Cells were precultivated under the following magnetic field conditions: NF: uniform magnetic field applied parallel to the oxygen gradient; SF: uniform magnetic field applied antiparallel to the oxygen gradient; GMF: ambient geomagnetic field (Bayreuth, Germany); ZF: ambient magnetic fields eliminated. Because Δ*mamAB* cells lack biomagnetism, swimming polarity is not selected during preculture, and in the hanging-drop assay cells do not preferentially accumulate at either drop edge facing magnetic north or south. The grey arrow indicates the magnetic field direction during the assay. The scale bar (50 µm) is shown in the top-left image and applies to all panels.

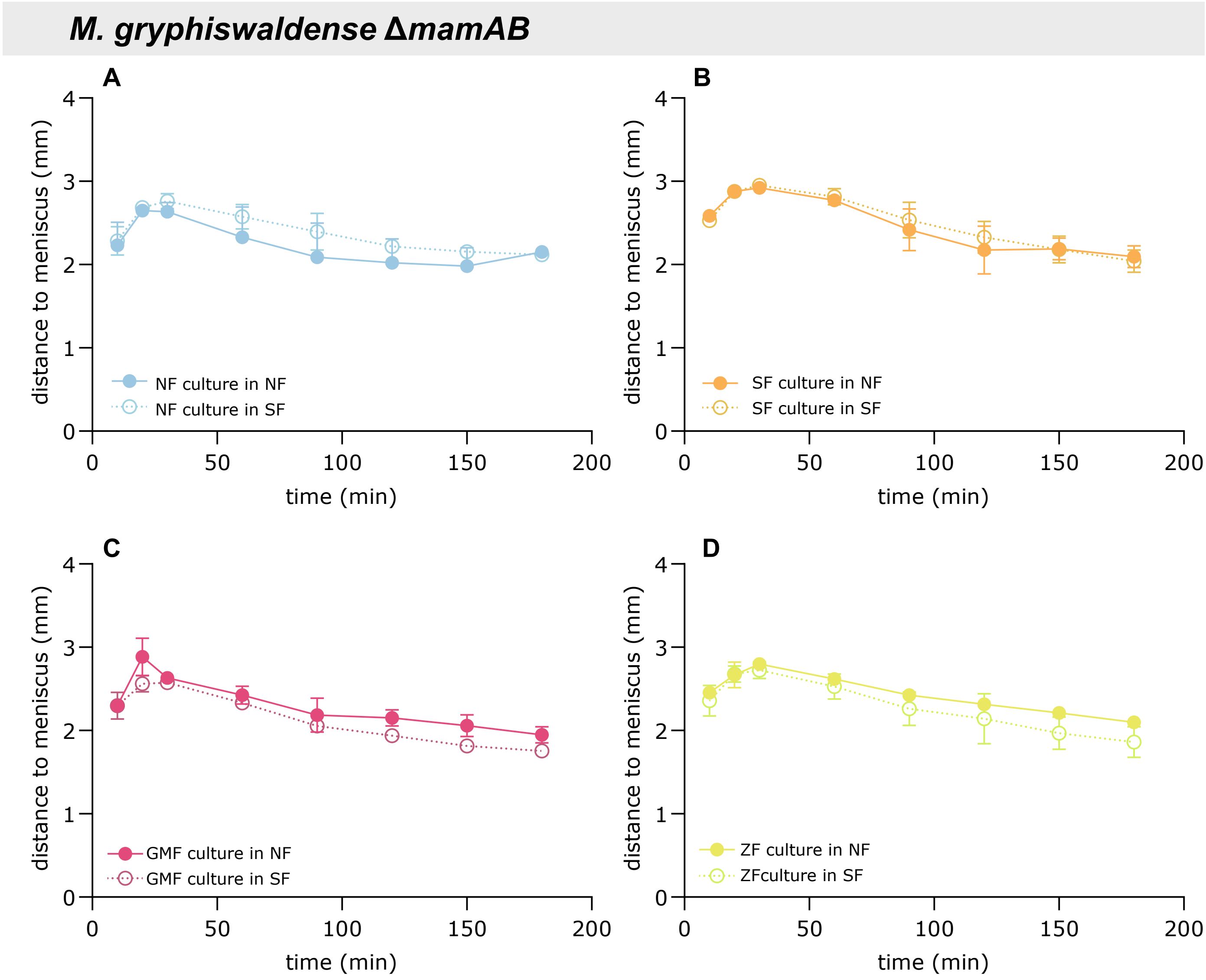


**Fig. S6. Spatio-temporal dynamics of aerotactic band formation for Δ*mamAB* strain cultures.** The distance between the aerotactic band and the air-liquid interface (meniscus) was measured at defined time points. **(A)** NF culture; **(B)** SF culture; **(C)** GMF culture; **(D)** ZF culture—each analyzed under NF and SF settings (solid and dashed lines, respectively). Data represent mean ± SD from $n$ = 3 microcapillary experiments and correspond to **Fig. 4**.

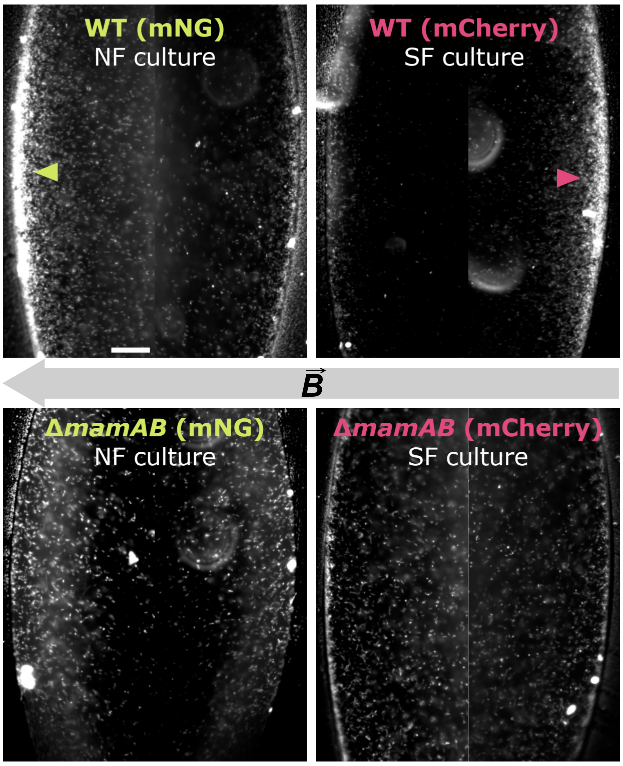


**Fig. S7. Hanging-drop assay of fluorescently labeled wild-type and Δ*mamAB* cells from NF and SF cultures.** mNG- and mCherry-labeled cells, precultivated under NF and SF conditions, respectively, were analyzed for swimming polarity in the hanging-drop assay before being mixed in equal amounts for competition assays. The grey arrow indicates the magnetic field direction. Dark-field micrographs are shown. The scale bar (50 µm) is depicted in the top-left image and applies to all panels. Data correspond to **Fig. 5**.

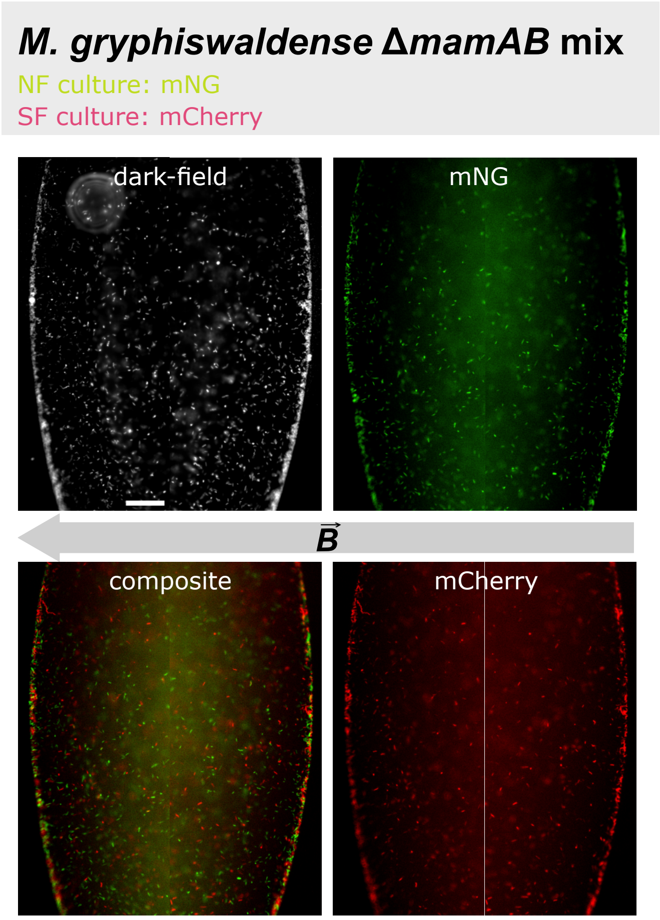


**Fig. S8. Hanging-drop assay of fluorescently labeled Δ*mamAB* cells.** mNG- and mCherry-labeled Δ*mamAB* cells, precultivated under NF and SF conditions, respectively, were analyzed in the hanging-drop assay after being mixed in equal amounts for competition assays. The grey arrow indicates the magnetic field direction. Dark-field micrographs are shown. The scale bar (50 μm) is depicted in the top-left image and applies to all panels.
