## Supplementary Tables for "Directional Matching of Swimming Polarity Provides a Competitive Advantage During Bacterial Magneto-Aerotaxis"

**Table S1. Strains and plasmids.** *E. coli* strains with plasmids are not individually listed.

| Strain or vector | Relevant characteristic(s) | Reference and/or source |
|---|---|---|
| **Strains** | | |
| *E. coli* | | |
| DH5α | Host for cloning; $F^-$ φ80*lacZ*ΔM15 Δ(*lacZYA-argF*)U169 *recA1 endA1 hsdR17*($r_K^-$ $m_K^+$) *phoA supE44* $\lambda^-$ *thi-1 gyrA96 relA1* | [1] |
| WM3064 | Conjugation strain; *thrB1004 pro thi rpsL hsdS lacZ*ΔM15 *RP4-1360* Δ(*araBAD*)*567* Δ*dapA1341*::[*erm pir*] | William Metcalf at UIUC |
| *M. gryphiswaldense* | | |
| MSR-1 R3/S1 | Laboratory wild type (WT); $Rif^R$, $Sm^R$ | [2] |
| Δ*mamAB* | *mamAB* operon deletion strain | [3] |
| MSR-1 R3/S1 Tn*7*:: $P_{mamDC45}$-*omNG100* | mNeonGreen-producing wild-type strain; $Km^R$ | This study |
| MSR-1 R3/S1 Tn*7*:: $P_{mamDC45}$-*mCherry* | mCherry-producing wild-type strain; $Km^R$ | This study |
| Δ*mamAB* Tn*7*:: $P_{mamDC45}$-*omNG100* | mNeonGreen-producing Δ*mamAB* strain; $Km^R$ | This study |
| Δ*mamAB* Tn7:: $P_{mamDC45}$-*mCherry* | mCherry-producing Δ*mamAB* strain; $Km^R$ | This study |
| **Plasmids** | | |
| pBAMII-Tn7 | Site-specific (*glmS* locus) Tn*7*-based insertion vector; *tnsABCD*, $Km^R$, $Amp^R$ | René Uebe at UBT (un-published) |
| pBAMII-$P_{mamDC45}$-*omNG100* | Vector for genomic insertion of $P_{mamDC45}$-*omNG100* | This study |
| pBAMII-$P_{mamDC45}$-*mCherry* | Vector for genomic insertion of $P_{mamDC45}$-*mCherry* | This study |

**Table S2. Oligonucleotides used for generating fluorescently labeled strains for competition experiments.** Restriction sites used for cloning are underlined.

| No. | Primer name | Sequence (5‘-3‘) |
|---|---|---|
| 558 | pBAMII-Tn7_fwd | GGAACTGCCTGGGCGAATTTAGC |
| 559 | pBAMII-Tn7_rev | AAATAGATGGGAACTGGGTGTAGCGTCG |
| 717 | PmamDC_SmaI/XmaI_fw2 | ACATCCCCGGGGCGAATTCCTCGAGCTTTTTCGCTT |
| 718 | omNG100_NotI_rev | CCGGCGGCCGCTCACTTATACAGTTCGTCCATGCCC |
| 719 | mCherry_NotI_rev | CCGGCGGCCGCTTACTTGTACAGCTCGTCCATGCC |
