## Supplementary material for "Directional Matching of Swimming Polarity Provides a Competitive Advantage During Bacterial Magneto-Aerotaxis": Legends for Supplementary Movies

**Movie S1: Aerotactic band shift of *M. gryphiswaldense* wild-type cells upon blue-light exposure.** Dark-field microscopy video (25 fps) at 200x magnification capturing the response of *M. gryphiswaldense* wild-type cells to blue light in a northern magnetic field (NF setting). The culture was grown under zero-field conditions, i.e., without conditions selecting for swimming polarity. Dark-field illumination was maintained throughout the recording. Between 5 s and 35 s, additional blue-light illumination was activated. The meniscus is positioned to the right. Yellow triangles mark the initial location of the aerotactic band; blue-light periods appear at bottom left. This movie is related to **Fig. 7**.

**Movie S2: Aerotactic band shift of *M. gryphiswaldense* Δ*mamAB* cells upon blue-light exposure.** Dark-field microscopy video (25 fps) at 200x magnification capturing the response of *M. gryphiswaldense* Δ*mamAB* cells to blue light in a northern magnetic field (NF setting). The aerotactic band comprises both mNG- and mCherry-labeled cells, as the video was recorded during competition experiments. Dark-field illumination was maintained throughout the recording. Between 5 s to 20 s, additional blue-light illumination was activated. The meniscus is positioned to the right. Yellow triangles mark the initial location of the aerotactic band; blue-light periods appear at bottom left.

**Movie S3: Green-light exposure does not induce an aerotactic band shift in *M. gryphiswaldense* Δ*mamAB* cells.** Dark-field microscopy video (25 fps) at 200x magnification revealed no detectable response upon activation of green light. However, a slight band shift can be observed when the green light was switched off. Dark-field illumination was maintained throughout the recording. Between 5 s to 20 s, additional green light was activated. The aerotactic band comprises both mNG- and mCherry-labeled Δ*mamAB* cells, however, an identical lack of response was also observed for wild-type cells (not shown). The capillary assay was performed under a northern field (NF setting). The meniscus is positioned to the right. Yellow triangles mark the initial location of the aerotactic band; green-light periods appear at bottom left.

**Movie S4: Long-term blue-light exposure.** Dark-field microscopy video (25 fps) at 200x magnification showing non-fluorescent wild-type cells precultured under zero-field conditions. Dark-field illumination was maintained throughout the recording. From 5 s onward, continuous blue-light illumination was applied for the entire duration (~12 min). The movie is shown as a time-lapse, accelerated by a factor of 16. During the recording, the microscope stage was adjusted to maintain the band within the visible region. The capillary assay was performed under a southern field (SF setting). The meniscus is positioned to the left. Blue-light periods appear at bottom left.
